# C_3_ gas-exchange analyses read bundle-sheath refixation in C_3_–C_4_ intermediate leaves as negative day respiration

**DOI:** 10.64898/2026.10.12.758566

**Authors:** Emre Yazar

## Abstract

C_3_–C_4_ intermediate (C_2_) plants decarboxylate photorespiratory glycine in the bundle sheath, where Rubisco refixes some of the released CO2, yet researchers analyze their gas exchange with methods built for C_3_ leaves. I calibrated the C_3_–C_4_ model of von Caemmerer against published compensation points and Laisk measurements on C_3_ and C_2_ *Flaveria* and relatives at 36 °C, and analyzed the resulting leaf as an experimenter would. With its true Rubisco kinetics and mesophyll conductance, Farquhar–von Caemmerer–Berry fits described its CO_2_ response closely but returned a day respiration (*R*_d_) of −5.4 µmol m^−2^ s^−1^ against a true 1.2, and underestimated maximum carboxylation and electron transport rates by 12% and 11%. The *R*_d_ deficit equaled net bundle-sheath CO_2_ fixation where mesophyll CO_2_ equals the photorespiratory compensation point (Γ*), within 5.3% in every leaf. The Laisk method gave Γ* = 42 µbar after mesophyll correction, compared with a true 65 µbar, and the oxygen-slope method overestimated Rubisco specificity by 26–53%. Published curves of two C_3_–C_4_ *Moricandia* species returned *R*_d_ of −2.5 µmol m^−2^ s^−1^, and their C_3_ congener 1.1. An *R*_d_ well below that of C_3_ relatives fitted the same way, signaling bundle-sheath refixation, not a fitting failure.

**Highlight:** In C_3_ fits of simulated C_3_–C_4_ intermediate leaves, day respiration falls below its true value due to net bundle-sheath CO_2_ fixation and turns negative; published *Moricandia* data agree.

## Introduction

In C_3_–C_4_ intermediate plants, photorespiratory glycine made in the mesophyll is decarboxylated in the bundle sheath, because glycine decarboxylase (GDC) in their leaves is confined to bundle-sheath mitochondria (Hylton *et al*., 1988; Rawsthorne *et al*., 1988; Rawsthorne, 1992). The CO_2_ released there raises the CO_2_ partial pressure around bundle- sheath Rubisco, which refixes some of it before it leaks back to the mesophyll (von Caemmerer, 1989; Monson and Rawsthorne, 2000). This photorespiratory CO_2_ pump, called C_2_ photosynthesis, lowers the CO_2_ compensation point well below that of C_3_ relatives (Holaday *et al*., 1984; Ku *et al*., 1991) and is regarded as a step on the route to C_4_ photosynthesis, which has evolved many times (Sage *et al*., 2011, 2012; Mallmann *et al*., 2014). Researchers study C2 species as models of this transition and as a source of traits to improve C_3_ crops (Bellasio and Farquhar, 2019; Lundgren, 2020).

Comparisons of C_3_–C_4_ intermediates with their C_3_ relatives rest largely on gas exchange (Ku *et al*., 1991; Vogan and Sage, 2011; Sage *et al*., 2013; Schlüter *et al*., 2017), and the tools for analyzing gas exchange were built for C_3_ leaves. *A*–*C*_i_ curves are fitted with the Farquhar– von Caemmerer–Berry (FvCB) model (Farquhar *et al*., 1980) to estimate the maximum Rubisco carboxylation rate (*V*_cmax_), the electron transport rate (*J*), and day respiration (*R*_d_) (Sharkey *et al*., 2007; Duursma, 2015; Busch *et al*., 2024). In the absence of day respiration, the CO2 compensation point (Γ*) and Rd are estimated from the common intersection of A–Ci lines measured at several low irradiances (the Laisk method; Laisk, 1977; Brooks and Farquhar, 1985; Walker and Ort, 2015). The in vivo CO_2_/O_2_ specificity of Rubisco, *S*_c/o_, can be obtained from how *C*_i_*, the *C*_i_ at which *A* = −*R*_d_, moves with O_2_ (Yin *et al*., 2020). In C_2_ species, the Laisk intersection, *C*\*, is used as a descriptor of how much photorespired CO_2_ the bundle sheath refixes (Sage *et al*., 2013).

Each of these tools treats the leaf as one C_3_ compartment, where all Rubisco sees the same CO_2_ partial pressure, and photorespired and respired CO_2_ is released only where that Rubisco can fix it. A C_2_ leaf differs on both counts. Its bundle-sheath Rubisco works at a higher CO_2_ partial pressure than its mesophyll Rubisco, and it refixes some of the CO2 released by GDC before it reaches the intercellular air spaces. The C_3_ equations have no term for either effect, so a fit must absorb both into the parameters the equations do contain. Even in C_3_ leaves, the location of (photo)respiratory CO_2_ release and its partial refixation complicate the interpretation of gas exchange (Tholen *et al*., 2012; Busch *et al*., 2013), and which C_3_ parameters can be trusted from which curves is still being worked out (Lochocki and McGrath, 2025; Xu *et al*., 2026). Yin *et al*. (2026) recently pointed out that applying the FvCB model as it stands to plants with synthetic photorespiratory bypasses is not advised, partly because bypasses change the amount and location of photorespiratory CO_2_ release. C_2_ photosynthesis is the natural case of such a change, yet how it biases each standard C_3_ analysis has not been worked out.

Here I ask what the standard C_3_ analyses return for a C_2_ leaf whose properties are known exactly. The leaf is the C_3_–C_4_ model of von Caemmerer (1989, 2000), which describes the mesophyll and bundle sheath, the glycine shuttle, and the CO_2_ leak from the bundle sheath explicitly. I calibrated it against the compensation points and Laisk measurements that Sage *et al*. (2013) made on C_2_ and C_3_ species of *Flaveria* and on C_3_ relatives at 36 °C, generated the measurements an experimenter would make, and analyzed them as an experimenter would: FvCB fits of *A*–*C*_i_ curves under seven common practices, the Laisk method, and the O_2_ method of Yin *et al*. (2020). The results follow a consistent pattern, which an identity derived from the C_3_ equations explains. I then test the main prediction, a negative fitted *R*_d_, on published *A*–*C*_i_ curves of C_3_ and C_3_–C_4_ species of *Moricandia* (Schlüter *et al*., 2017).

## Materials and methods

### Leaf model

Symbols are defined in Supplementary Table S1. The C_2_ leaf is the C_3_–C_4_ model of von Caemmerer (1989), in the form summarized by von Caemmerer (2000, 2013), with no C_4_ cycle (phosphoenolpyruvate carboxylation set to zero), as in intermediates without significant C_4_-cycle activity (Sage *et al*., 2012). The leaf has two photosynthetic compartments, mesophyll (subscript m) and bundle sheath (subscript s), each with its own Rubisco and electron transport. Net CO_2_ assimilation is

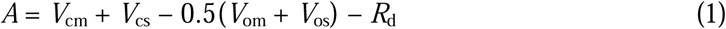

where *V*_c_ and *V*_o_ are the Rubisco carboxylation and oxygenation rates, and *R*_d_ is day respiration, split equally between the compartments (*R*_m_ = *R*_s_ = *R*_d_/2; Alonso-Cantabrana and von Caemmerer, 2016). CO_2_ reaches the mesophyll chloroplasts through the mesophyll conductance *g*_m_,

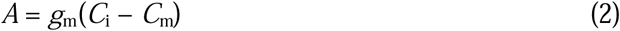

and the mesophyll balance is

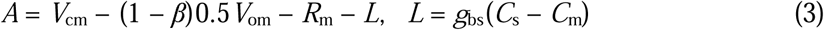

where *L* is the leak of CO_2_ from the bundle sheath to the mesophyll through the bundle- sheath conductance *g*_bs_, and β is the fraction of mesophyll glycine decarboxylated in the bundle sheath (β = 1 when GDC is confined to the bundle sheath; Hylton *et al*., 1988). The rest of the bundle sheath’s CO_2_ supply is its own photorespiration and *R*_s_. In each compartment *k*, Rubisco is limited by its own carboxylation capacity or by its own electron transport:

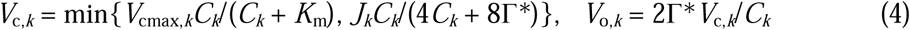

with *K*_m_ = *K*_c_(1 + *O*/*K*_o_). A fraction *f*_s_ of the leaf’s Rubisco and electron transport sits in the bundle sheath (*V*_cmax,s_ = *f*_s_*V*_cmax_, *J*_s_ = *f*_s_*J*), so a C_3_ leaf and a C_2_ leaf with the same *V*_cmax_ and *J* carry the same capacity per unit leaf area. Whole-leaf *J* follows a non-rectangular hyperbola in the incident photosynthetic photon flux density (PPFD, µmol photons m^−2^ s^−1^), *I*,

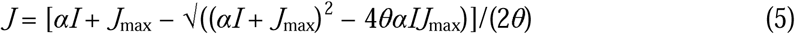

with initial slope α and curvature θ = 0.7. The C_3_ leaf is the special case *f*_s_ = 0, β = 0; in that limit the implementation reproduced an independent solution of the C_3_ FvCB model to within 3 × 10^−11^ µmol m^−2^ s^−1^. For a given *C*_m_, the bundle-sheath balance has a single root in *C*_s_, so the model was solved as two nested one-dimensional root problems with Brent’s method.

A second variant tested the explanation Sage *et al*. (2013) offered for the fall of the C_2_ compensation point with irradiance: residual GDC in the mesophyll decarboxylates glycine up to a capacity *D*_m_, and glycine produced in excess of it overflows to the bundle sheath, so the effective β rises with the photorespiratory flux.

CO_2_ is expressed as partial pressure (µbar) and O_2_ in mbar. The published data are mole fractions (µmol mol^−1^), which I treat as numerically equal to partial pressures at the measurement pressure (about 1 bar).

### *Calibration against Sage et al.* (2013)

The C_3_ control was calibrated against the C_3_ group of Sage *et al*. (2013), measured at 36 °C (Table 1): the Laisk intersection *C*\* (60.0 ± 0.6 µmol mol^−1^, mean ± SE of sunflower, *Sartwellia flaveriae* and *F. cronquistii*; their Table III), the *C*_i_ at which the high-light *A*–*C*_i_ response crosses *A* = −*R*_d_ (60.6 ± 0.6; Table III), the apparent *R*_d_ of the Laisk analysis (mean of the two C_3_ panels of their Fig. 8, 1.28 µmol m^−2^ s^−1^), and Γ at six irradiances (their Fig. 7, digitized; means of *S. flaveriae* and *F. cronquistii*; Supplementary Table S2). On the model leaf I ran their procedure as reconstructed from their Methods and Fig. 8: straight lines fitted to seven *A*–*C*_i_ points between 50 and 100 µbar at PPFDs of 100, 150, 200, 300 and 400 µmol m^−2^ s^−1^; *C*\* and the apparent *R*_d_ (*R*_d,app_) taken as the least-squares common point of the five lines; Γ at each irradiance as the *C*_i_ intercept of its line; and the high-light *C*_i_ at *R*_d_ from a line through seven points from Γ − 5 to Γ + 30 µbar at 1200 µmol m^−2^ s^−1^ (their high light was 1000–1400), extrapolated to *A* = −*R*_d,app_. Group targets were the mean of the two species’ plotted values, with an uncertainty of half their difference but no less than 2 µmol mol^−1^; *R*_d,app_ uncertainty was set to 0.15 (C_3_) and 0.30 (C_2_) µmol m^−2^ s^−1^ because Fig. 8 shows single representative leaves. Free parameters were Γ* at 25 °C, *R*_d_, α, and *V*_cmax_. Fixed were *g*_m_ (0.3 mol m^−2^ s^−1^ bar^−1^), *J*_max_ (450 µmol m^−2^ s^−1^) and the Michaelis constants of Rubisco from *F. floridana* (*K*_c_ 395 µbar, *K*_o_ 544 mbar at 25 °C; Kubien *et al*., 2008, as tabulated by Alonso- Cantabrana and von Caemmerer, 2016). Both leaves had this Rubisco, so any differences between them reflect the C_2_ mechanism; in vitro, the CO_2_/O_2_ specificity of Rubisco from C_3_– C_4_ *Flaveria* does not differ from that of its C_3_ congeners (Perdomo *et al*., 2015). Kinetic constants were scaled to 36 °C with the activation energies of Bernacchi *et al*. (2001); *V*_cmax_, *J*_max,_ and *R*_d_ were specified directly at 36 °C. Parameters were minimized by the sum of squared standardized residuals (χ^2^) with a trust-region least-squares algorithm.

**Table 1.** Calibration of the C_3_ control and the C_2_ leaves against Sage *et al*. (2013) at 36 °C.

|  | C<sub>3</sub> target | C<sub>3</sub> control | C<sub>2</sub> target | C<sub>2</sub> leaf (best fit) | C<sub>2</sub> overflow |
| --- | --- | --- | --- | --- | --- |
| *Calibration targets and model values* |  |  |  |  |  |
| C* (μbar) | 60.0 ± 0.6 | 60.4 | 37.0 ± 1.4 | 37.2 | 37.5 |
| High-light C<sub>i</sub> at R<sub>d</sub> (μbar) | 60.6 ± 0.6 | 60.2 | 22.5 ± 1.5 | 21.0 | 20.9 |
| R<sub>d,app</sub> (μmol m<sup>-2</sup> s<sup>-1</sup>) | 1.28 ± 0.15 | 1.28 | 1.48 ± 0.30 | 1.57 | 1.55 |
| Γ at PPFD 100 (μbar) | 100.2 ± 7.2 | 97.9 | 103.1 ± 6.1 | 84.5 | 88.2 |
| Γ at PPFD 150 (μbar) | 87.0 ± 4.0 | 87.0 | 69.8 ± 2.4 | 71.5 | 71.4 |
| Γ at PPFD 200 (μbar) | 80.8 ± 3.5 | 81.6 | 65.0 ± 2.6 | 64.5 | 63.8 |
| Γ at PPFD 300 (μbar) | 75.9 ± 3.3 | 76.2 | 55.8 ± 3.0 | 57.2 | 56.8 |
| Γ at PPFD 400<sup>b</sup> (μbar) | 73.4 ± 3.0 | 73.5 | 50.7 ± 3.1 | 53.5 | 53.6 |
| Γ at high light<sup>c</sup> (μbar) | 71.5 ± 2.0 | 71.4 | 33.6 ± 2.0 | 35.4 | 35.1 |
| χ<sup>2</sup> (d.f.) |  | 1.2 (5) |  | 12.8 (6) | 9.5 (5) |
| *Parameters at 36 °C* |  |  |  |  |  |
| Γ* (μbar; 25 °C in parentheses) |  | 64.6 (37.5) |  | 64.6 (37.5) | 64.6 (37.5) |
| K<sub>m</sub> (μbar) |  | 1518 |  | 1518 | 1518 |
| V<sub>cmax</sub> (μmol m<sup>-2</sup> s<sup>-1</sup>) |  | 298 |  | 298 | 298 |
| J<sub>max</sub> (μmol m<sup>-2</sup> s<sup>-1</sup>) |  | 450<sup>a</sup> |  | 450<sup>a</sup> | 450<sup>a</sup> |
| α (mol mol<sup>-1</sup>) |  | 0.345 |  | 0.345 | 0.345 |
| g<sub>m</sub> (mol m<sup>-2</sup> s<sup>-1</sup> bar<sup>-1</sup>) |  | 0.3<sup>a</sup> |  | 0.3<sup>a</sup> | 0.3<sup>a</sup> |
| R<sub>d</sub> (μmol m<sup>-2</sup> s<sup>-1</sup>) |  | 1.27 |  | 1.24 | 1.17 |
| f<sub>s</sub> |  | 0<sup>a</sup> |  | 0.125 | 0.104 |
| g<sub>bs</sub> (mmol m<sup>-2</sup> s<sup>-1</sup> bar<sup>-1</sup>) |  | — |  | 10.8 | 2.7 |
| β |  | 0<sup>a</sup> |  | 1<sup>a</sup> | variable<sup>d</sup> |
| D<sub>m</sub> (μmol m<sup>-2</sup> s<sup>-1</sup>) |  | — |  | — | 2.23 |

The C_2_ leaf kept every parameter of the C_3_ control except three: *f*_s_, *g*_bs_ and *R*_d_, fitted with β = 1 to the C_2_ group of Sage *et al*. (2013) (*F. sonorensis* and *F. angustifolia*; *C*\* 37.0 ± 1.4 and high-light *C*_i_ at *R*_d_ 22.5 ± 1.5 µmol mol^−1^, Table III; *R*_d,app_ 1.48 µmol m^−2^ s^−1^ from their Fig. 8; Γ at six irradiances from their Fig. 7) with the Laisk window moved to 35–100 µbar, as in their C_2_ panels. Setting β = 1 idealizes these species; Sage et al. (2013) found some GDC in the mesophyll, and the overflow variant represents this residual mesophyll activity. The fit was started from a grid of initial values. Because *g*_bs_ proved poorly constrained, I profiled χ^2^ along it, refitting *f*_s_ and *R*_d_ at 13 fixed values of *g*_bs_. Besides the best fit, three leaves from this profile (*g*_bs_ = 20, 2 and 0.1 mmol m^−2^ s^−1^ bar^−1^) and the overflow variant, with *D*_m_ as a fourth free parameter, were carried through every analysis, giving five C_2_ leaves in all.

### Temperature

Every analysis of the synthetic measurements was repeated at 25 °C, a common measurement temperature at which photorespiration is lower. *V*_cmax_, *J*_max_, *R*_d_ and *D*_m_ were scaled with the activation energies of Bernacchi *et al*. (2001, 2003), *D*_m_ with that of *R*_d_, without high-temperature deactivation, and the kinetic constants as above; *g*_m_, *f*_s_, *g*_bs_ and β were held constant. The 25 °C leaves are therefore extrapolations of the 36 °C calibration, not calibrations in their own right.

### Synthetic measurements and their analysis

*A*–*C*_i_ curves were generated at a PPFD of 1500 µmol m^−2^ s^−1^ at 14 values of *C*_i_ from 40 to 1400 µbar (40, 60, 80, 100, 130, 170, 220, 300, 400, 550, 700, 900, 1100 and 1400). They were fitted with the C_3_ FvCB model,

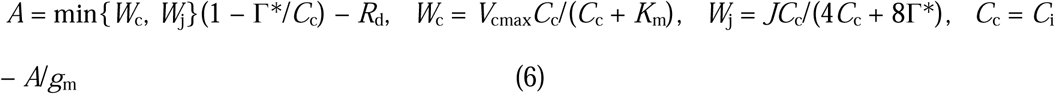

under seven practices that differ in what is taken as known (Table 2). P1 uses the true Γ*, *K*_m_ and *g*_m_. P2 uses the true Γ* and *K*_m_ with infinite *g*_m_, i.e., a *C*_i_-based fit. P3 uses the constants of Bernacchi *et al*. (2001) in a *C*_i_-based fit, the default of widely used fitting software (Duursma, 2015). P4 fits Γ* with the true *K*_m_ and *g*_m_, and P5 fits Γ* with the Bernacchi *K*_m_ in a *C*_i_-based fit. P6 and P7 fix *R*_d_ at the Laisk value and use the true Γ* (P6) or fit it (P7). “True” values are those of the generating leaf, which are also those of the C_3_ control; *V*_cmax_, *J,* and, where not fixed, *R*_d_ were free in every practice. The minimum in Equation 6 was taken over the two gross carboxylation rates: below Γ* a minimum over net rates selects the wrong limitation, and variants of this kind are known to change parameter estimates (Lochocki and McGrath, 2025). Each fit was started from every split of the curve into a Rubisco-limited lower part and an electron-transport-limited upper part, and the best solution was refined by least squares on the full model. Parameter bounds (*V*_cmax_ and *J* 1–3000 µmol m^−2^ s^−1^, *R*_d_ −10 to 20 µmol m^−2^ s^−1^, Γ* 0–300 µbar) were reached by no fit.

**Table 2.** What standard C_3_ analyses return for the C_3_ control and the best C_2_ leaf.

| Analysis | $\Gamma^*$, $K_m$, $g_m$<br>used | $C_3$ control | | | $C_2$ leaf (best fit) | | |
| --- | --- | --- | --- | --- | --- | --- | --- |
| | | $V_{\text{cmax}}$<br>ratio | $R_d$ | $\Gamma^*$ | $V_{\text{cmax}}$<br>ratio | $R_d$ | $\Gamma^*$ [RMSE] |
| 36 °C |  |  |  |  |  |  |  |
| True values |  | 1 | 1.27 | 64.6 | 1 | 1.24 | 64.6 |
| P1 ($A-C_i$ fit) | true, true, true | 1.00 | 1.27 | – | 0.88 | –5.39 | – [0.13] |
| P2 ($A-C_i$ fit) | true, true, ∞ | 0.69 | 1.18 | – | 0.64 | –3.13 | – [0.81] |
| P3 ($A-C_i$ fit) | B01, B01, ∞ | 0.79 | –0.08 | – | 0.74 | –4.30 | – [0.61] |
| P4 ($A-C_i$ fit) | fitted, true, true | 1.00 | 1.27 | 64.6 | 0.88 | –5.97 | 67.8 [0.08] |
| P5 ($A-C_i$ fit) | fitted, B01, ∞ | 0.78 | –4.29 | 107.9 | 0.72 | –8.60 | 111.4 [0.22] |
| P6 ($A-C_i$ fit) | true, true, true | 1.00 | 1.28<sup>a</sup> | – | 1.95 | 1.57<sup>a</sup> | – [2.46] |
| P7 ($A-C_i$ fit) | fitted, true, true | 1.00 | 1.28<sup>a</sup> | 64.5 | 0.92 | 1.57<sup>a</sup> | 30.9 [1.16] |
| Laisk | –, –, true | – | 1.28 | 64.7<br>(60.4) | – | 1.57 | 42.4 (37.2) |
| O<sub>2</sub> slope | –, –, – | – | true | 64.7 | – | true | 49.0 |
| 25 °C |  |  |  |  |  |  |  |
| True values |  | 1 | 0.65 | 37.5 | 1 | 0.64 | 37.5 |
| P1 ($A-C_i$ fit) | true, true, true | 1.00 | 0.65 | – | 0.88 | –3.57 | – [0.18] |
| P2 ($A-C_i$ fit) | true, true, ∞ | 0.77 | 1.56 | – | 0.71 | –1.22 | – [0.71] |
| P3 ($A-C_i$ fit) | B01, B01, ∞ | 0.90 | 0.42 | – | 0.83 | –2.29 | – [0.50] |
| P4 ($A-C_i$ fit) | fitted, true, true | 1.00 | 0.65 | 37.5 | 0.87 | –4.42 | 42.2 [0.08] |
| P5 ($A-C_i$ fit) | fitted, B01, ∞ | 0.88 | –1.47 | 56.4 | 0.81 | –4.90 | 63.2 [0.31] |
| P6 ($A-C_i$ fit) | true, true, true | 1.00 | 0.67<sup>a</sup> | – | 1.25 | 1.14<sup>a</sup> | – [1.36] |
| P7 ($A-C_i$ fit) | fitted, true, true | 1.00 | 0.67<sup>a</sup> | 37.5 | 0.97 | 1.14<sup>a</sup> | 19.9 [0.83] |
| Laisk | –, –, true | – | 0.67 | 37.5<br>(35.3) | – | 1.14 | 21.1 (17.2) |
| O<sub>2</sub> slope | –, –, – | – | true | 37.7 | – | true | 25.8 |

The Laisk analysis was run as in the calibration, with its *C*_i_ windows scaled with Γ* at 25 °C. The O_2_ method of Yin *et al*. (2020) was applied with their design: *A*–*C*_i_ curves at a PPFD of 1000 µmol m^−2^ s^−1^ at 20, 100, 210, 350 and 500 mbar O_2_, and *C*_i_*, the *C*_i_ intercept of a line through five points of *A* + *R*_d_ against *C*_i_ near *A* = −*R*_d_, with *R*_d_ taken as known. In a C_3_ leaf,

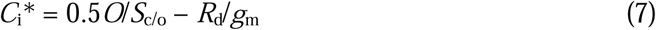

provided that (photo)respired CO_2_ is released where Rubisco works (the classical case, *m* = 0 in Yin *et al*., 2020), so the slope of *C*_i_* against *O* is 0.5/*S*_c/o_, and Γ* = 0.5*O*/*S*_c/o_. The C_3_ control satisfies this assumption by construction. I also computed the slopes over 20–210 and 210–500 mbar separately.

### Respiration deficit and net bundle-sheath fixation

For every C_2_ leaf, I computed the net CO_2_ fixation by bundle-sheath Rubisco, *N*_s_ = *V*_cs_ − 0.5*V*_os_, at the *C*_i_ where the mesophyll CO_2_ partial pressure *C*_m_ equals Γ*, and compared it with the deficit *R*_d_ − *R*_d,fit_ returned by practice P1. Leaf-level carboxylation-to-oxygenation ratios, (*V*_cm_ + *V*_cs_)/(*V*_om_ + *V*_os_), relative to the C_3_ control were computed at *C*_i_ of 150, 200 and 250 µbar and a PPFD of 1200 µmol m^−2^ s^−1^ for comparison with Keerberg *et al*. (2014).

### Calibration sensitivity

*J*_max_ and *g*_m_ cannot be identified from the calibration data. The whole calibration and analysis were therefore repeated with *g*_m_ of 0.2 and 0.5 mol m^−2^ s^−1^ bar^−1^ and with *J*_max_ of 300 and 800 µmol m^−2^ s^−1^ (Supplementary Table S3). *V*_cmax_ was only loosely constrained by the calibration targets: the five calibrations placed it between 226 and 414 µmol m^−2^ s^−1^.

### Measurement noise

Each experiment (one *A*–*C*_i_ curve, the Laisk set and the O_2_ set) was repeated 200 times for the C_3_ control and the best C_2_ leaf, with independent Gaussian noise added to every value of *A* (standard deviation 0.3 µmol m^−2^ s^−1^, and 0.6 in a second set), and the analysis was rerun on each replicate: practices P1, P3 and P4, the Laisk analysis and the O_2_ method, at 36 °C and, with the lower noise, at 25 °C.

### Published data

Representative *A*–*C*_i_ curves of the C_3_ *Moricandia moricandioides* and the C_3_–C_4_ *M. suffruticosa* and *M. arvensis* (line MOR1) were digitized from Fig. 2A of Schlüter *et al*. (2017; CC BY 4.0), measured at 25 °C and a PPFD of 1500 µmol m^−2^ s^−1^. The page was rendered at 600 dpi from the published PDF; the axes were calibrated from the tick marks, and each marker was located by template matching, which separates the overlapping markers at low *C*_i_ (Supplementary Fig. S1). The procedure recovered 13 points per species (Supplementary Table S4). Medians of Γ and of the initial *A*–*C*_i_ slope (8–10 plants per line; their Fig. 2B, C) were read from the same rendered page by locating the median line of each box. No Rubisco kinetic constants are available for *Moricandia*, so each curve was fitted under five settings: P3 and *C*_c_-based fits with tobacco constants (*K*_c_ 260 µbar, *K*_o_ 179 mbar and Γ* = 38.1 µbar from *k*_c_ 3.6 and *k*_o_ 0.9 s^−1^; von Caemmerer *et al*., 2009, as tabulated by Johnson and Berry, 2021) at *g*_m_ of 0.2, 0.3 and 0.5 mol m^−2^ s^−1^ bar^−1^, and with Γ* free at *g*_m_ = 0.3.

### Statistical analysis

The simulated leaves are deterministic, so there is no sampling variance to test. The calibration fits report χ^2^ against the standardized published targets, and the profile along *g*_bs_ shows which parameters those targets constrain. The Monte Carlo replicates give the spread an experimenter would see from one curve per leaf; they are summarized as mean ± standard deviation, the fraction of replicates with a negative fitted *R*_d_, and the 5th, 50th and 95th percentiles of the RMSE and of r^2^. For the published data, with one digitized curve per *Moricandia* line, inference rests on agreement between the single curves and the population medians, not on statistical tests.

### Software and use of AI

All computations used Python 3.13.16 with NumPy 2.5.3, SciPy 1.18.1 and Matplotlib 3.11.2. One script runs the entire analysis and writes a single results file; every number in this paper comes from that file or the deposited data files, and we deposited the code, the digitized data, and the results file (Data availability). A large language model, Claude (Anthropic; Opus 5.5, model ID claude-opus-5-5), was used under the author’s direction to help check the code, the published figures, and the text edits. It was not used as a source of scientific claims. The author checked the derivations, analyses, citations, and conclusions against the code, the result file, and the primary literature, and takes full responsibility for the content.

## Results

### The calibrated leaves reproduce the C_3_ and C_2_ phenotypes

The C_3_ control reproduced every C_3_ target (χ^2^ = 1.2 on 5 degrees of freedom, d.f.; Fig. 1A; Table 1). It gave *C*\* = 60.4 µbar against 60.0 ± 0.6 µmol mol^−1^, a high-light *C*_i_ at *R*_d_ of 60.2 against 60.6 ± 0.6, and a fall of Γ from 97.9 µbar at a PPFD of 100 µmol m^−2^ s^−1^ to 71.4 at high light. Its fitted Γ* is 37.5 µbar at 25 °C and 64.6 µbar at 36 °C, with *R*_d_ = 1.27 µmol m^−2^ s^−1^.

**Fig. 1.**
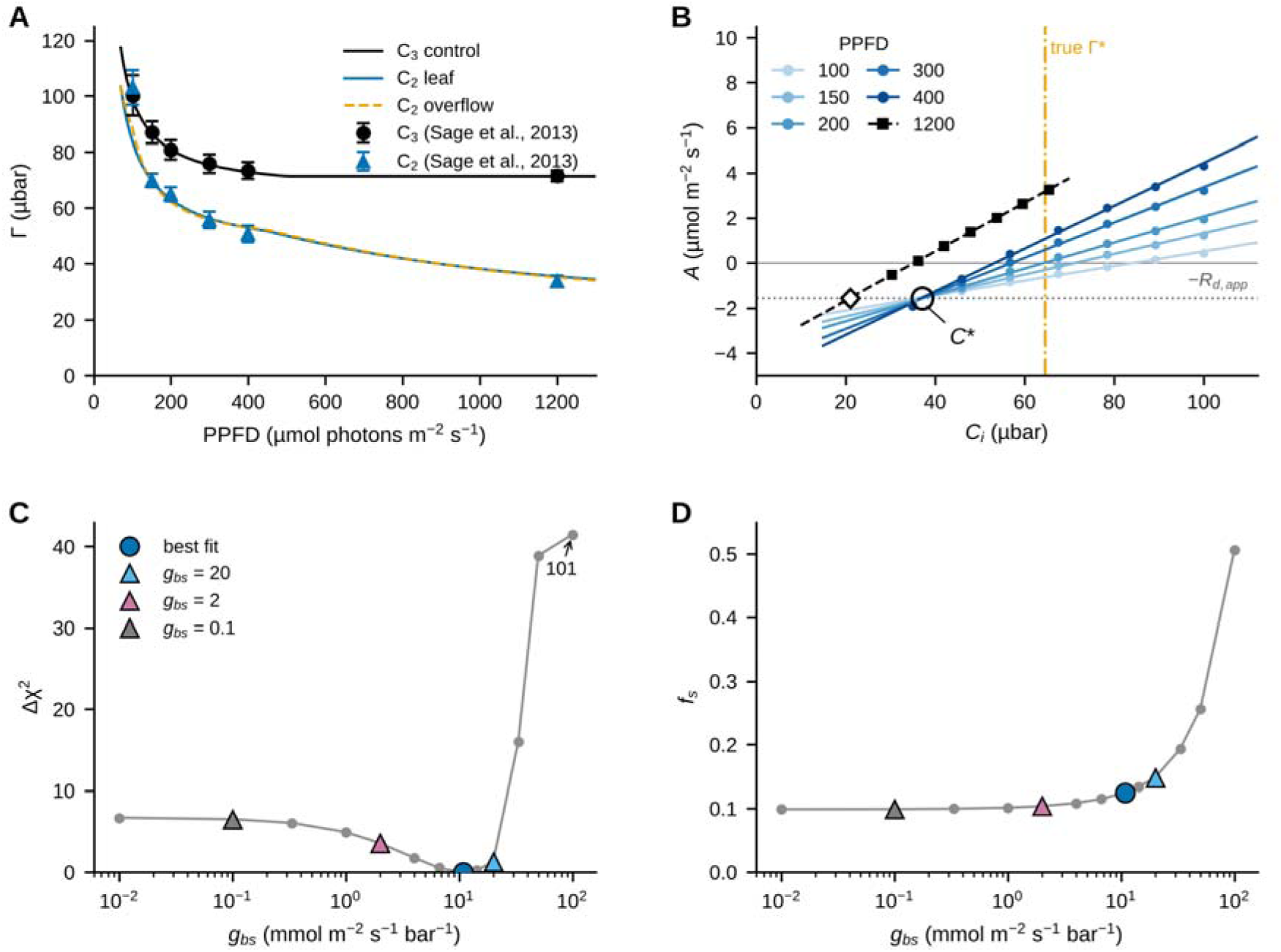
Calibration of the model leaves against Sage *et al*. (2013) at 36 °C. (A) CO_2_ compensation point Γ against PPFD: group means of the C_3_ species (*Sartwellia flaveriae* and *F. cronquistii*; circles) and the C_2_ species (*F. sonorensis* and *F. angustifolia*; triangles) digitized from their Fig. 7, with the uncertainties used in the fit; high-light values (measured at 1000–1400 µmol m^−2^ s^−1^) are plotted at 1200; lines, compensation points of the C_3_ control, the best C_2_ leaf and the overflow variant. (B) The Laisk procedure run on the C_2_ leaf: *A*–*C*_i_ points and fitted lines at PPFDs of 100–400 µmol m^−2^ s^−1^ (light to dark blue) and the high- light line at 1200 µmol m^−2^ s^−1^ (squares, dashed line). The open circle marks the intersection (*C*\*, −*R*_d,app_; dotted line), the open diamond the high-light *C*_i_ at *R*_d_, and the dash-dotted line the true Γ* of the leaf’s Rubisco. (C) Increase in χ^2^ along the bundle-sheath conductance *g*_bs_, with *f*_s_ and *R*_d_ refitted at each value, and (D) the corresponding *f*_s_. Large circle, best fit; triangles, the three further leaves carried through every analysis. The off-scale point in (C) is labeled with its value. Alt text (Fig. 1). Four panels. Panel A plots the compensation point against light. Black circles for the C_3_ group fall from about 100 to 72 µbar and level off, with a black line following them; blue triangles for the C_2_ group fall from about 103 to 34, with a blue line following them except at the lowest light, where it lies about 19 µbar below the point. Panel B shows five straight lines of increasing slope that converge near a *C*_i_ of 37 and an *A* of −1.6, well to the left of a vertical line at the true Γ* near 65; a steeper dashed line crosses the dotted horizontal line near a *C*_i_ of 21. Panel C plots the increase in χ^2^ against conductance on a logarithmic axis: it is near zero between about 5 and 20, rises steeply above 20 (to 39 at 50 and 101 at 100), and rises slowly to about 7 as conductance falls to 0.01. Panel D shows the bundle-sheath fraction staying between 0.10 and 0.15 below a conductance of 20 and rising steeply to 0.5 at 100.

**Fig. 2.**
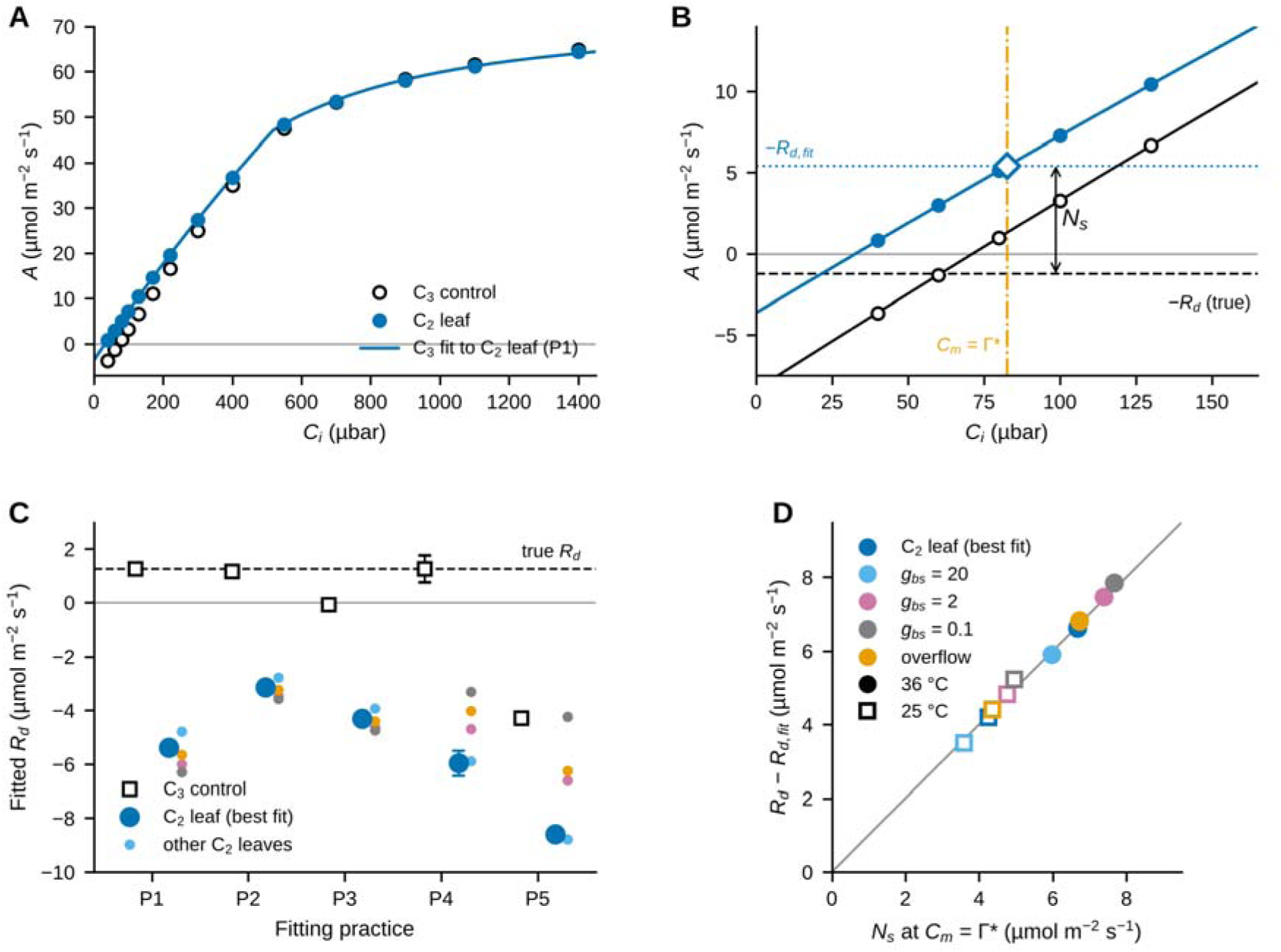
C_3_ fits of the C_2_ leaf at 36 °C. (A) *A*–*C*_i_ curves of the C_3_ control (open circles) and the C_2_ leaf (closed circles) at a PPFD of 1500 µmol m^−2^ s^−1^, with the C_3_ FvCB fit to the C_2_ leaf under P1 (true Γ*, *K*_m_ and *g*_m_; line). (B) Detail at low *C*_i_. The C_3_ control (black line) reaches −*R*_d_ (dashed line), where its chloroplast CO_2_ partial pressure equals Γ*. The C_2_ leaf, at the *C*_i_ where its mesophyll CO_2_ partial pressure equals Γ* (dash-dotted line), is higher than −*R*_d_ by *N*_s_, the net fixation by bundle-sheath Rubisco; the C_3_ fit (blue line) passes through this point (diamond) and therefore returns −*R*_d,fit_ = *N*_s_ − *R*_d_ (dotted line). (C) Fitted *R*_d_ under practices P1–P5 for the C_3_ control (squares), the best C_2_ leaf (large circles) and the other four C_2_ leaves (small circles); error bars, standard deviation over 200 noisy replicates (σ = 0.3 µmol m^−2^ s^−1^; P1, P3 and P4 only), mostly smaller than the symbols; dashed line, true *R*_d_. (D) The respiration deficit *R*_d_ − *R*_d,fit_ (P1) against *N*_s_ for the five C_2_ leaves at 36 °C (closed symbols) and 25 °C (open symbols); line, 1:1. Alt text (Fig. 2). Four panels. Panel A shows two nearly coincident curves rising from about zero to 64 µmol m^−2^ s^−1^ over a *C*_i_ of 0–1400 µbar; the fitted blue line passes through the closed circles of the C_2_ leaf, and the open circles of the C_3_ control lie slightly lower at low and intermediate *C*_i_. Panel B enlarges the low-*C*_i_ region: both sets of points lie on straight lines, the C_2_ line several units above the C_3_ line. A vertical dash-dotted line meets the C_2_ line at a diamond at about 5.4; a dotted horizontal line at that height marks minus the fitted *R*_d_, a dashed line at −1.2 marks minus the true *R*_d_, and a double arrow between them is labeled *N*_s_. Panel C plots fitted *R*_d_ for five practices: squares for the C_3_ control lie near the true *R*_d_ for P1, P2 and P4, near zero for P3 and at −4.3 for P5; circles for the C_2_ leaves lie between −2.8 and −8.8 for every practice. Panel D shows ten points for the C_2_ leaves on the 1:1 line between about 3.6 and 7.7 µmol m^−2^ s^−1^.

The C_2_ leaf maintained the same Rubisco, electron transport, and mesophyll conductance, and allocated 12.5% of its Rubisco and electron transport to the bundle sheath, with a conductance of 10.8 mmol m−2 s−1 bar−1 and *R*_d_ = 1.24 µmol m^−2^ s^−1^ (χ^2^ = 12.8 on 6 d.f.). It reproduced *C*\* (37.2 against 37.0 ± 1.4), the high-light *C*_i_ at *R*_d_ (21.0 against 22.5 ± 1.5; Fig. 1B) and Γ from a PPFD of 150 µmol m^−2^ s^−1^ upward (Fig. 1A). Its one clear failure is Γ at the lowest irradiance, 84.5 µbar against 103.1 measured (standardized residual −3.0): the measured C_2_ compensation point rises more steeply toward low light than the model’s. The overflow variant, in which the bundle sheath receives a growing share of glycine as irradiance rises, improved the fit only modestly (χ^2^ = 9.5 with one more parameter; Γ at 100 µmol m^−2^ s^−1^ 88.2 µbar).

The bundle-sheath conductance was poorly constrained (Fig. 1C, D). Taking an increase in χ^2^ of 3.84 × χ_min_^2^/d.f. = 8.2 as an approximate 95% threshold, which allows for the misfit of the best fit, the data excluded leaky bundle sheaths (*g*_bs_ above about 25 mmol m^−2^ s^−1^ bar^−1^; χ^2^ rose by 39 at 50 and by 101 at 100) but not tight ones: as *g*_bs_ fell from the optimum to 0.01, χ^2^ rose by only 6.7. Without the allowance for misfit (a threshold of 3.84), values below about 1.7 would also be excluded. Over the range of *g*_bs_ from 20 to 0.1 mmol m^−2^ s^−1^ bar^−1^, *f*_s_ stayed between 0.10 and 0.15. The data therefore fix how much Rubisco works in the bundle sheath more firmly than they fix the barrier around it. Results below are given for the best C_2_ leaf and, as ranges, for all five C_2_ leaves.

### C_3_ fits describe C_2_ A–C_i_ curves closely but return a negative R_d_

Fitted with the true Γ*, *K*_m_ and *g*_m_ (P1), the C_3_ model recovered the C_3_ control exactly and described the curve of the C_2_ leaf with a root-mean-square error (RMSE) of 0.13 µmol m^−2^ s^−1^ (Fig. 2A; 0.13–0.60 across the five C_2_ leaves; Supplementary Fig. S2). However, the parameters were wrong (Table 2). *R*_d_ came out at −5.39 µmol m^−2^ s^−1^ against a true 1.24, and *V*_cmax_ and *J* were underestimated by 12% and 11%. The underestimates follow from the negative *R*_d_: the C_3_ model subtracts *R*_d_ from every point of the curve, so a negative value raises the whole modeled curve, and the capacities must fall to keep it on the data.

Every practice that left *R*_d_ free returned a negative value for every C_2_ leaf, from −8.8 to −2.8 µmol m^−2^ s^−1^ at 36 °C and from −4.9 to −1.1 at 25 °C (Fig. 2C; Supplementary Table S5). The sign alone, however, is not diagnostic under every practice. Under P1 and P4, the C_3_ control returned its true *R*_d_ and under P2 a positive one, but under the common practice P3 it was near zero at 36 °C (−0.08 µmol m^−2^ s^−1^; negative in 76% of noisy replicates), because the C_3_ control’s compensation point (71.4 µbar) lies just below the Bernacchi Γ* (73.6 µbar). P5 gave the C_3_ control clearly negative values (−4.29 at 36 °C and −1.47 at 25 °C), because a free Γ* in a *C*_i_-based fit absorbs the CO_2_ drawdown across the mesophyll. What distinguished the C_2_ leaf was the size of the deficit: under P3 its *R*_d_ was lower than the C_3_ control’s by 4.2 µmol m^−2^ s^−1^ at 36 °C and by 2.7 at 25 °C.

### The respiration deficit equals net bundle-sheath fixation

The deficit has a simple origin (Fig. 2B). At the *C*_i_ where the mesophyll CO_2_ partial pressure equals Γ*, mesophyll Rubisco gains nothing because its carboxylation is exactly balanced by the CO_2_ that its oxygenation releases. Net assimilation there is the bundle-sheath gain minus respiration, *A* = *N*_s_ − *R*_d_. The C_3_ model, fitted with the true Γ* and *g*_m_, assigns that point the same chloroplast CO_2_ partial pressure, because *C*_c_ = *C*_i_ − *A*/*g*_m_ is computed from the same *A*, and at *C*_c_ = Γ* it predicts *A* = −*R*_d,fit_. A fit that passes through the curve at that point must therefore return

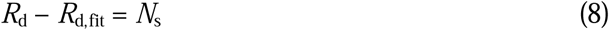

(derivation in Supplementary Methods S1). Across the five C_2_ leaves at both temperatures, the deficit matched *N*_s_ within 5.3% (Fig. 2D); what remains is the misfit of the C_3_ curve at that single point. *N*_s_ was 6.0–7.7 µmol m^−2^ s^−1^ at 36 °C and 3.6–5.0 at 25 °C, reached in the best C_2_ leaf at *C*_i_ ≈ 83 and 50 µbar.

A similar argument applies to *C*_i_-based fits, which place Γ* on the *C*_i_ axis: if such a fit passed through the curve at *C*_i_ = Γ*, −*R*_d,fit_ would equal the observed *A* there. Where the initial part of the curve is nearly linear, with slope *s* and *C*_i_ intercept Γ,

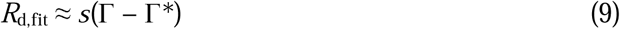

so a leaf whose compensation point lies well below the Γ* used in the fit returns a negative *R*_d_. This needs only Γ and the initial slope, both routinely reported for C_3_–C_4_ intermediates. Equation 9 is approximate because *C*_i_-based fits do not pass exactly through the curve at Γ*: for the simulated leaves, the fitted *R*_d_ exceeded −*A*(*C*_i_ = Γ*) by 0.13–1.17 µmol m^−2^ s^−1^ under P2 and P3. Its sign was right in 23 of the 24 fits; the exception was the C_3_ control at 25 °C under P3, whose Γ (40.8 µbar) lies only 1.9 µbar below the Bernacchi Γ*: Equation 9 gives −0.23 µmol m^−2^ s^−1^, the fit 0.42.

### *Freeing* Γ*\* or fixing R_d_ moves the error rather than removing it*

Letting Γ* float did not rescue *R*_d_. With the true *g*_m_ (P4), the fitted Γ* ranged from 48.2 to

70.7 µbar across the C_2_ leaves (true 64.6), and *R*_d_ stayed between −5.97 and −3.30 µmol m^−2^ s^−1^. A single *A*–*C*_i_ curve does not contain the information needed to separate refixation from Rubisco specificity. Forcing *R*_d_ to the Laisk value moved the error elsewhere. With Γ* fixed (P6), *V*_cmax_ rose to 1.9–2.0 times its true value, and the fit failed visibly (RMSE 2.3–2.7 µmol m^−2^ s^−1^). With Γ* free (P7), the fit placed Γ* at 24.5–33.6 µbar, about half the true value, at an RMSE of 0.68–1.16. An analyst who constrains *R*_d_ to positive values therefore obtains either a poor fit or a low Γ*, and a low Γ* would be read as a more CO_2_-specific Rubisco.

### *The Laisk intersection of a C_2_ leaf is not* Γ*

The Laisk lines of the C_2_ leaf converged, as they did in the measurements, at *C*\* = 37.2 µbar (Fig. 1B), far below the true Γ* of 64.6 µbar. The standard correction for mesophyll conductance, Γ* = *C*\* + *R*_d,app_/*g*_m_ (von Caemmerer, 2000), recovered Γ* in the C_3_ control (64.7 µbar), as it must because the control releases (photo)respired CO_2_ where Rubisco works, but returned 41.4–44.9 µbar for the C_2_ leaves (Fig. 3A). The apparent *R*_d_ at the intersection was also too high: 1.57 against a true 1.24 µmol m^−2^ s^−1^ (+26%). At 25 °C, *C*\* was 17.2 µbar and the corrected value was 21.1 µbar, compared with Γ* = 37.5 µbar (Fig. 3B). The intersection reflects the whole leaf, set by how much photorespired CO_2_ the bundle sheath recaptures, not by its Rubisco.

**Fig. 3.**
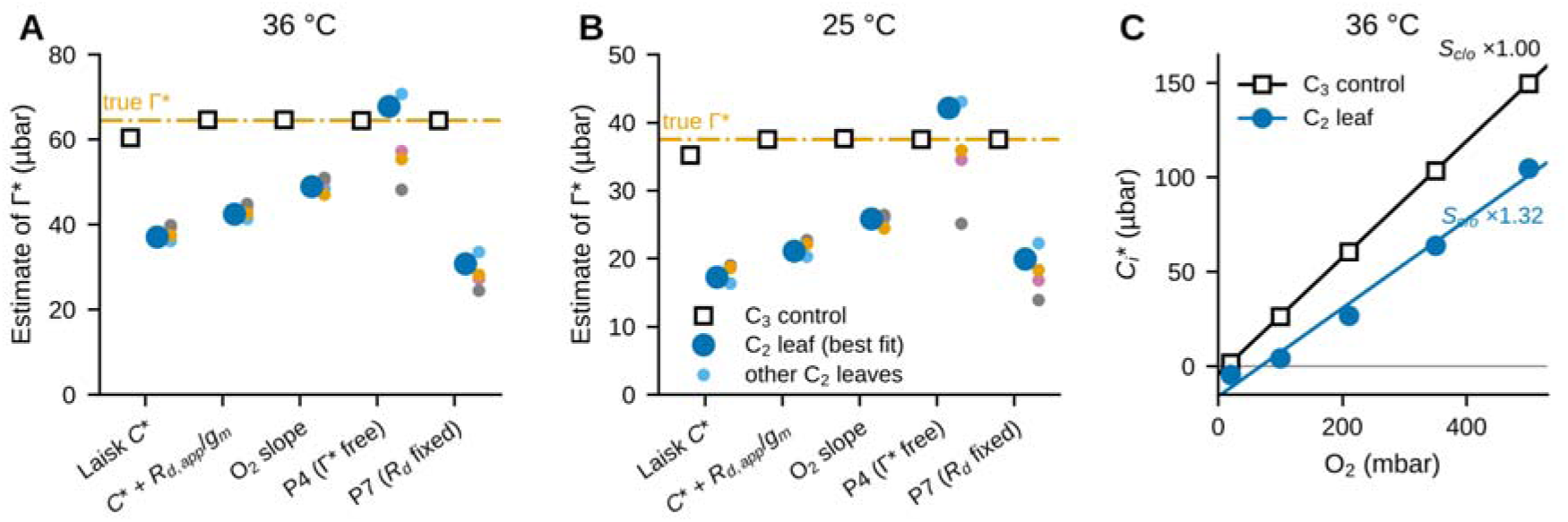
Estimates of Γ* and of Rubisco specificity. (A, B) Γ* estimated by five methods for the C_3_ control (squares), the best C_2_ leaf (large circles) and the other four C_2_ leaves (small circles) at 36 °C (A) and 25 °C (B): the Laisk intersection *C*\*; *C*\* corrected for mesophyll conductance (*C*\* + *R*_d,app_/*g*_m_); 0.5*O*/*S*_c/o_ from the O_2_ slope; and Γ* fitted to the *A*–*C*_i_ curve with *R*_d_ free (P4) or fixed at the Laisk value (P7). Dash-dotted line, true Γ*. (C) *C*_i_* against O_2_ at 36 °C in the design of Yin *et al*. (2020) for the C_3_ control and the best C_2_ leaf, with regression lines; labels give *S*_c/o_ from the slope relative to its true value. Alt text (Fig. 3). Three panels. Panels A and B are dot plots with five method categories on the horizontal axis. The squares of the C_3_ control lie on or just below a dash-dotted line at the true Γ* (65 µbar in A, 37.5 in B). The circles of the C_2_ leaves lie well below that line for the Laisk, corrected Laisk, O_2_-slope, and P7 methods (24–51 µbar in A and 14–26 in B) and near it for P4, except for the leaf with the tightest bundle sheath (48.2 µbar in A and 25.2 in B). Panel C plots *C*_i_* against O_2_ from 0 to 500 mbar: the squares of the C_3_ control lie on a line rising from about 2 to 150 µbar, and the circles of the C_2_ leaf on a shallower line from about −5 to 105; the labels read 1.00 and 1.32 times the true specificity.

### The O_2_ slope overestimates Rubisco specificity

In the C_3_ control, which meets the assumption behind Equation 7 by construction, *C*_i_* rose linearly with O_2_ and its slope returned *S*_c/o_ within 1% (Fig. 3C). In the C_2_ leaves at 36 °C, *C*_i_* also rose almost linearly (r^2^ = 0.981–0.987; 0.975–0.989 at 25 °C), but too slowly, so the five-point slope overestimated *S*_c/o_ by a factor of 1.26–1.37 at 36 °C and 1.42–1.53 at 25 °C. The relation was not straight: at 36 °C the slope between 20 and 210 mbar O_2_ gave 1.72–2.07 times the true *S*_c/o_ and that between 210 and 500 mbar 1.14–1.15 times (1.92–2.48 and 1.24– 1.26 at 25 °C). At 20 mbar O_2_ and 36 °C, *A* of the C_2_ leaves stayed above −*R*_d_ at every positive *C*_i_, because the bundle sheath refixes part of the CO_2_ released by its own mitochondria; the line-based *C*_i_* was negative. The Γ* implied by the O_2_ slope at 36 °C was 47.1–51.1 µbar against a true 64.6 µbar.

### Measurement noise does not hide the signatures

With noise of standard deviation 0.3 µmol m^−2^ s^−1^, P1 fits of the C_2_ leaf were as good as those of the C_3_ control (median RMSE 0.30 against 0.25 µmol m^−2^ s^−1^; Supplementary Table S6), so goodness of fit gave no warning. The negative *R*_d_, however, lay far outside the noise: P1 returned −5.41 ± 0.19 µmol m^−2^ s^−1^ for the C_2_ leaf and 1.28 ± 0.22 for the C_3_ control. At 36 °C, every C_2_ replicate gave a negative *R*_d_ under P1, also at twice the noise, and no C_3_ replicate did. *C*\* scattered more (37.4 ± 4.0 µbar) but stayed far below Γ*. The O_2_ method returned *S*_c/o_ = 2.15 ± 0.05 mbar µbar^−1^ against a true 1.63 (at 25 °C, 4.08 ± 0.17 against 2.80), with a median r^2^ of 0.985.

### Calibration choices change magnitudes, not conclusions

Repeating the calibration with a *g*_m_ of 0.2 or 0.5 mol m^−2^ s^−1^ bar^−1^ or a *J*_max_ of 300 or 800 µmol m^−2^ s^−1^ changed *f*_s_ to 0.114–0.126 and the P1 *R*_d_ of the C_2_ leaf to between −8.2 and −3.5 µmol m^−2^ s^−1^, but not the pattern (Supplementary Table S3). *V*_cmax_ was 0.87–0.91 times its true value, the corrected Laisk value was 40–43 µbar against a Γ* of 63–67, the O_2_ method overestimated *S*_c/o_ by a factor of 1.32–1.42, and Equation 8 held within 6% in every calibration.

### Published C_3_–C_4_ Moricandia curves return a negative R_d_

The central prediction can be tested on measurements: a C_3_–C_4_ leaf fitted as a C_3_ leaf should return a negative *R*_d_, and its C_3_ relative a positive one. Under P3, the digitized curves of Schlüter *et al*. (2017) gave *R*_d_ = 1.14 µmol m^−2^ s^−1^ for the C_3_ *M. moricandioides*, and −2.48 and −2.49 for the C_3_–C_4_ *M. suffruticosa* and *M. arvensis*, with RMSE of 0.41, 0.35 and 0.57 µmol m^−2^ s^−1^ (Fig. 4; Supplementary Table S7). The sign held under every setting of the kinetic constants. With tobacco constants and a *g*_m_ of 0.2–0.5 mol m^−2^ s^−1^ bar^−1^, *R*_d_ was 2.7 to 3.5 for the C_3_ species and −4.3 to −2.3 for the C_3_–C_4_ lines. With Γ* free, the C_3_ species placed Γ* at 30.6 µbar with *R*_d_ = 5.42, whereas the C_3_–C_4_ lines placed it at 47.9 and 47.8 µbar, above the tobacco value, and *R*_d_ fell further (−4.61 and −4.60), as for the best C_2_ leaf. The representative curves are single leaves, but the population medians tell the same story: Equation 9, with the median Γ and initial slope of 8–10 plants per line, gives *R*_d_ ≈ 1.32 µmol m^−2^ s^−1^ for *M. moricandioides* and −2.25 and −1.38 for the C_3_–C_4_ lines (Fig. 4B), margins well above the misfit of the fits (RMSE 0.35–0.57). The C_3_–C_4_ minus C_3_ difference under P3, −3.6 µmol m^−2^ s^−1^, is similar in size to the difference the model predicts for the *Flaveria* pair at 25 °C (−2.7).

**Fig. 4.**
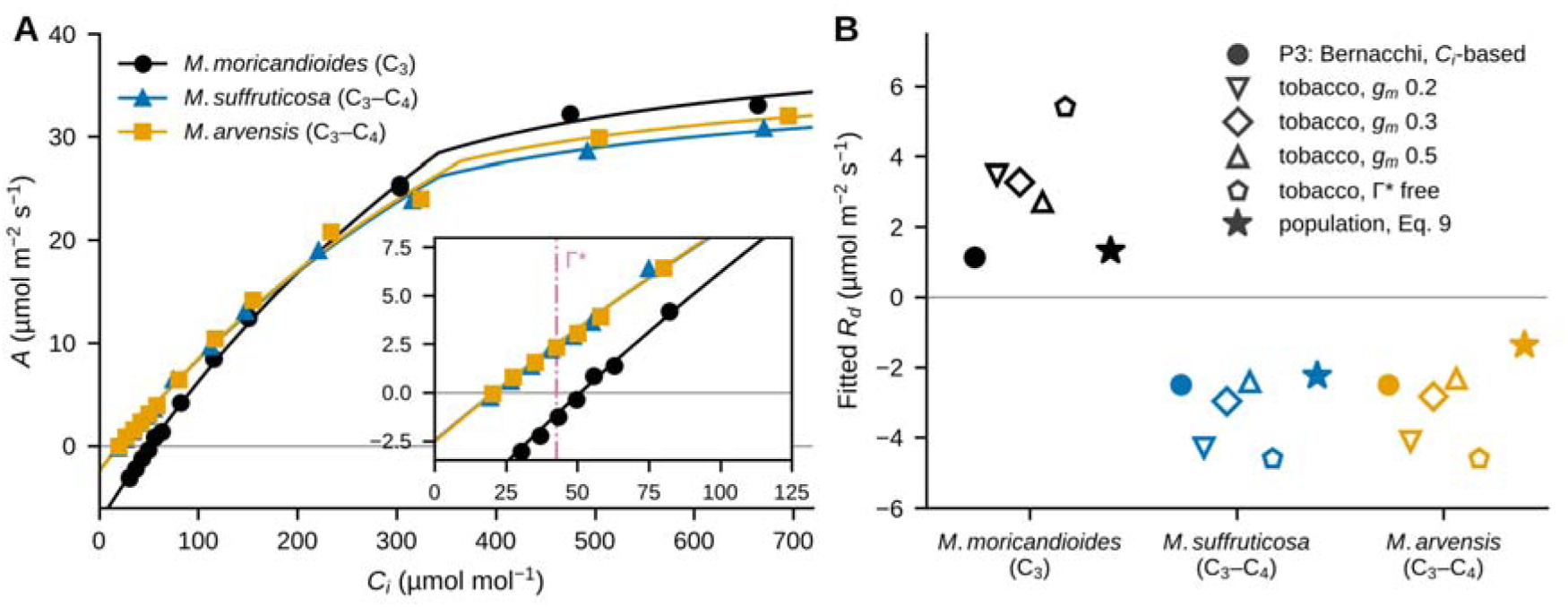
C_3_ fits of published *A*–*C*_i_ curves of *Moricandia* (digitized from Schlüter *et al*., 2017, Fig. 2A; CC BY 4.0). (A) The C_3_ *M. moricandioides* (circles) and the C_3_–C_4_ *M. suffruticosa* (triangles) and *M. arvensis* MOR1 (squares), with C_3_ FvCB fits under P3 (constants of Bernacchi *et al*., 2001, *C*_i_-based; lines). Inset: low *C*_i_, with Γ* = 42.75 µmol mol^−1^ (dash-dotted line). The C_3_–C_4_ curves are above zero at Γ*, so their fits return a negative *R*_d_. (B) Fitted *R*_d_ for each species under five settings of the kinetic constants (P3, closed symbols; tobacco constants at three values of *g*_m_ and with Γ* fitted, open symbols) and from Equation 9 with the median Γ and initial slope of 8–10 plants per line (stars). Alt text (Fig. 4). Two panels. Panel A shows *A*–*C*_i_ points for three species rising from about −3 to 33 µmol m^−2^ s^−1^ over a *C*_i_ of 20–700 µmol mol^−1^, with fitted lines; at low *C*_i_ the two C_3_–C_4_ species lie on nearly identical curves above that of the C_3_ species. An inset of *C*_i_ from 0 to 125 shows the C_3_–C_4_ lines crossing zero near 22 and passing about 2.5 units above zero at a vertical line at Γ*, while the C_3_ line crosses zero near 51. Panel B plots fitted *R*_d_ for the three species: every symbol for the C_3_ species is positive (1.1 to 5.4), and every symbol for the two C_3_–C_4_ species is negative (−1.4 to −4.6).

## Discussion

### A negative fitted R_d_ measures refixation

A negative *R*_d_ from an *A*–*C*_i_ fit is easily taken as a sign that something went wrong, such as too few points at low *C*_i_, an error in *C*_i_, or a poorly conditioned fit. For C_3_–C_4_ intermediates, the results here say the opposite. A C_3_ fit that leaves *R*_d_ free and uses the true Γ* must return a negative *R*_d_ for a C_2_ leaf whose net bundle-sheath fixation, where mesophyll CO_2_ equals Γ*, exceeds its respiration, because its *A*–*C*_i_ curve is already above zero where the C_3_ model places the zero of mesophyll net carboxylation (Equation 8); under *C*_i_-based practices the same holds approximately whenever Γ lies well below the Γ* used (Equation 9). The deficit is not noise. It equals that net bundle-sheath fixation within a few percent in every simulated leaf, and it survives measurement noise, calibration choices, and temperature. Where *R*_d_ is known independently, a C_3_ fit therefore measures net bundle-sheath fixation at that point: *N*_s_ = *R*_d_ − *R*_d,fit_. *N*_s_ is not the leaf’s total refixation, but it is a direct, quantitative readout of the C_2_ pump. In the simulated leaves, the Laisk estimate, although itself shifted, served well enough for this purpose: *R*_d,app_ − *R*_d,fit_ overestimated *N*_s_ by at most 14.1% across all leaves and both temperatures. A better independent estimate of *R*_d_ would tighten this estimate of *N*_s_.

The same reasoning explains the published curves. The C_3_–C_4_ *Moricandia* lines have compensation points of 19–25 µmol mol^−1^ (population medians), about half the C_3_ value of Γ* used by default (42.75 µmol mol^−1^), so Equation 9 requires them to return a negative *R*_d_, by margins (1.4–2.3 µmol m^−2^ s^−1^) well above the misfit of the fits. The single curves and the population medians agree, and the C_3_–C_4_ minus C_3_ difference under the default practice is similar in size to that predicted for the *Flaveria* model at the same temperature. Taken together, an *R*_d_ well below that of C_3_ relatives fitted the same way is a quantitative signature of the glycine shuttle, not an artifact to be removed by constraining the fit.

### What the other parameters mean

Because the negative *R*_d_ is a constant offset, it is paid for by the capacities. In C_3_ fits of C_2_ leaves, *V*_cmax_ was 0.87–0.91 and *J* 0.84–0.92 times their true values across all calibrations. A C_2_ species with exactly the Rubisco and electron transport capacity of its C_3_ relative would therefore appear to have about a tenth less of both. The bias is modest, but it has a known sign, and it bears on any comparison of photosynthetic capacity or nitrogen-use efficiency between C_3_ and C_3_–C_4_ relatives that rests on C_3_ fits.

Freeing Γ* did not help, and constraining *R*_d_ to positive values made matters worse. With *R*_d_ fixed, the curve could only be fitted by halving Γ* (P7), and such a fit would be read as evidence for a more CO_2_-specific Rubisco; with Γ* also fixed (P6), the fit failed visibly, a failure that could easily be blamed on the measurements. For the best C_2_ leaf, the unconstrained fits gave no warning, because their residuals were as small as those of a C_3_ leaf; for the other C_2_ leaves the misfit was larger (RMSE up to 0.60 µmol m^−2^ s^−1^ without noise) but lay mostly at intermediate and high *C*_i_, away from the low-*C*_i_ region that decides *R*_d_ (Supplementary Fig. S2). Goodness of fit, the usual check on an FvCB analysis (Sharkey *et al*., 2007; Busch *et al*., 2024), need not detect a missing compartment; only a comparison with C_3_ relatives under the same practice, or an analysis that includes the bundle sheath, can.

### Laisk and oxygen methods

Sage *et al*. (2013) used *C*\* as an indicator of photorespiratory refixation in C_2_ species, and the results support that use. *C*\* depends on how much photorespired CO_2_ the bundle sheath recaptures. What *C*\* cannot be is Γ*. Even with the correction for mesophyll conductance, the intersection of a C_2_ leaf lay 31–46% below the Γ* of its own Rubisco. Using a C_2_ *C*\* as Γ* in an FvCB analysis, or comparing it with in vitro specificities, conflates a leaf-level concentrating effect with an enzyme property. The uncertainty in Γ* is already large in C_3_leaves (Walker *et al*., 2017); in C_2_ leaves, the Laisk estimate is also biased.

Yin et al.’s (2020) O_2_ method inherits the same problem. Bundle-sheath refixation makes *C*_i_* rise more slowly with O_2_ than the specificity of Rubisco implies, and the relation stays close enough to linear that r^2^ gives no warning. A C_2_ leaf therefore appears to carry a Rubisco about a quarter to a half more specific than it does. In vitro, Rubisco of C_3_–C_4_ *Flaveria* has the specificity of its C_3_ congeners (Perdomo *et al*., 2015), so an in vivo *S*_c/o_ above the C_3_ value in a C_3_–C_4_ species is expected from refixation alone and is not evidence for a different enzyme.

### Limitations

The model leaf is a type I intermediate with no C_4_ cycle, calibrated on two *Flaveria* species at one temperature, with a common bundle-sheath fraction for Rubisco and electron transport and equal respiration in the two compartments. Three discrepancies with measurements point to what it may miss. First, it does not reproduce the steep rise of the C_2_ compensation point at the lowest irradiance, and neither does the overflow mechanism Sage et al. (2013) proposed; the low-light behavior of C_2_ needs another explanation or another measurement. Second, the model’s leaf-level ratio of carboxylation to oxygenation was only 1.03–1.24 times that of the C_3_ control, whereas Keerberg *et al*. (2014) inferred from ^14^CO_2_ labeling a 3.2-fold higher mean CO_2_ at Rubisco in *F. pubescens* than in C_3_ *F. cronquistii*. *Flaveria pubescens* has some C_4_-cycle activity; the conditions differ, and the calibration species are not the same, but the gap is large. If real C_2_ leaves concentrate CO_2_ more strongly than the model does, *N*_s_, and with it the bias in *R*_d_, would be larger. Third, the 25 °C results are extrapolations with *g*_m_, *f*_s_ and *g*_bs_ held constant. For the published curves, the digitized data are single representative leaves, and no kinetic constants exist for *Moricandia* Rubisco; the conclusions about them rest on the sign of *R*_d_, which was the same under every setting tried and agreed with the population medians.

The sign of the effect does not depend on these details. Equations 8 and 9 need only that the curve be above zero where the C_3_ model places Γ*, that is, that the leaf’s compensation point lie below the C_3_ value of Γ*. That is the defining phenotype of C_3_–C_4_ intermediates.

### Recommendations

Three practical recommendations follow. First, fit C_3_–C_4_ intermediates without bounding *R*_d_ at zero, report the fitted *R*_d_, and interpret its difference from that of C_3_ relatives fitted the same way as a measure of net bundle-sheath fixation (Equation 8). Second, report *C*\* and Γ as descriptors of the C_2_ phenotype, as Sage *et al*. (2013) did, and do not use them as Γ*; take Rubisco kinetics of C_2_ species from in vitro measurements (Kubien *et al*., 2008; Perdomo *et al*., 2015). Third, when Rubisco and electron transport capacities of C_2_ leaves matter, fit a model that contains the bundle sheath (von Caemmerer, 2000), with *f*_s_ and *g*_bs_ constrained by data a single *A*–*C*_i_ curve cannot supply: Γ at several irradiances constrained *f*_s_ here but not *g*_bs_, which needs further information such as carbon isotope discrimination (Alonso-Cantabrana and von Caemmerer, 2016).

## Supplementary data

The following supplementary data are available at JXB online.

**Fig. S1.** Digitization check: the panel of Schlüter *et al*. (2017, Fig. 2A; CC BY 4.0) with the marker positions found by template matching.

**Fig. S2.** Practice-P1 fits and their residuals for the C_3_ control and the five C_2_ leaves at 36 and 25 °C.

**Table S1.** Symbols, definitions and units.

**Table S2.** Digitized compensation points of *Flaveria* and *Sartwellia* (Sage *et al*., 2013, Fig. 7).

**Table S3.** Calibration sensitivity: calibrations with *g*_m_ of 0.2, 0.3 and 0.5 mol m^−2^ s^−1^ bar^−1^ and *J*_max_ of 300, 450 and 800 µmol m^−2^ s^−1^, and what each analysis returns for each.

**Table S4.** Digitized *A*–*C*_i_ data of *Moricandia* (Schlüter *et al*., 2017, Fig. 2A).

**Table S5.** All fitting practices for all six leaves at 36 and 25 °C.

**Table S6.** Monte Carlo summary for the C_3_ control and the best C_2_ leaf at two noise levels (36 °C) and at 25 °C.

**Table S7.** Fits of the digitized *Moricandia* curves under five settings of the kinetic constants, with the digitized population medians.

**Methods S1.** Derivation of Equations 8 and 9.

## Acknowledgements

The use of a large language model is described in Materials and Methods.

## Author contributions

EY: conceptualization; methodology; software; formal analysis; investigation; data curation; visualization; writing – original draft; writing – review and editing.

## Data availability

All code, digitized data, and the single result file from which every number in this paper is read are openly available at Zenodo (https://doi.org/10.5281/zenodo.23319739). One script reproduces the whole analysis, and an audit script checks every quantitative statement in the text against the result file. No new experimental data were generated; the measurements analyzed are published in Sage *et al*. (2013) and Schlüter *et al*. (2017).

## Abbreviations

*A*: net CO_2_ assimilation rate
*C*\*: intercellular CO_2_ partial pressure at the common intersection of Laisk lines
*C*_i_: intercellular CO_2_ partial pressure
*C*_i_*: *C*_i_ at which *A* = −*R*_d_
*C*_m_ and *C*_s_: CO_2_ partial pressures in mesophyll and bundle-sheath chloroplasts
FvCB: Farquhar–von Caemmerer–Berry
GDC: glycine decarboxylase
*g*_bs_: bundle-sheath conductance to CO_2_
*g*_m_: mesophyll conductance
Γ: CO_2_ compensation point
Γ*: CO_2_ compensation point in the absence of day respiration
*J*: electron transport rate
*N*_s_: net CO_2_ fixation by bundle-sheath Rubisco
PPFD: photosynthetic photon flux density
*R*_d_: day respiration
RMSE: root-mean-square error
*S*_c/o_: Rubisco CO_2_/O_2_ specificity
*V*_cmax_: maximum Rubisco carboxylation rate.

